# X chromosome association analyses using multiple models identifies 18 genetic loci influencing dietary intake in UK Biobank

**DOI:** 10.64898/2026.04.24.720538

**Authors:** Maizy S Brasher, Kristen J Sutton, William B Patterson, Joanne B Cole

**Affiliations:** Department of Biomedical Informatics, School of Medicine, University of Colorado Anschutz, Aurora, CO

## Abstract

Although dietary intake is a leading risk factor for many common diseases, adherence to dietary recommendations remains low. This may partly reflect limited consideration of individual differences in eating behavior that arise from both environmental and genetic factors. While genome-wide association studies (GWAS) of dietary intake have identified hundreds of associated loci, the X chromosome has largely been ignored. To address this gap, we applied multiple X-chromosome-wide association study (X-WAS) models on dietary intake phenotypes to identify novel associations.

We performed X-WAS of 46 dietary intake traits from food frequency questionnaires in up to 424,758 European participants from the UK Biobank. Phenotypes included quantitative measures (e.g., fruit intake), binary traits (e.g., decaffeinated vs caffeinated coffee), and principal component-derived food groups. We tested for genetic associations using several models: a traditional sex-combined additive GWAS, additive models stratified by sex, and two joint models accounting for sex-interaction effects and non-additivity. We also conducted X-WAS in five additional genetic ancestry groups and performed a sex-combined multi-ancestry additive GWAS meta-analysis with up to 445,773 individuals.

We identified 18 loci associated with 20 dietary intake traits (*P* < 5×10^-8^), including 17 variants without prior associations in the GWAS Catalog. Among these loci, 10 were significant across multiple X-WAS models, and 5 were strongest in a model other than the traditional sex-combined additive GWAS, highlighting the value of approaches that address known complexities of the X chromosome.

These results demonstrate that incorporating the X chromosome in GWAS can reveal novel loci, even for complex behavioral traits such as dietary intake. Applying multiple association models further improves discovery by accounting for unique features of the X chromosome.

**Author Summary:** Although diet is a major risk factor for many common diseases, adherence to healthy eating guidelines remains low. One reason is that current recommendations do not account for individual differences in food choice that arise from environmental or genetic factors. Previous genetic studies have identified hundreds of genetic variants associated with dietary behaviors, but most have excluded the X chromosome due to its analytical complexity and differences between males and females. However, accumulating evidence suggests that the X chromosome contains important genetic variation that impacts complex traits.

We analyzed data from hundreds of thousands of individuals to identify genetic variants on the X chromosome associated with dietary intake. To address the unique features of the X chromosome, we applied multiple different models that account for sex-differences and non-additive genetic effects. We identified 18 regions in the genome associated with at least one dietary intake trait. These results reveal new insights into the genetics underlying eating behavior and highlight the importance of incorporating the X chromosome in genetic studies of complex traits.

## Introduction

Dietary intake is a leading risk factor for many common, noncommunicable diseases, including type 2 diabetes, obesity, and cardiovascular disease [1,2]. The 2017 Global Burden of Diseases, Injuries, and Risk Factors Study attributed 11 million annual deaths to dietary risk factors, particularly high sodium intake, low intake of whole grains, and low intake of fruits [1]. Despite the known detrimental effects of an unhealthful diet, adherence to recommendations remains poor [1], in part because current guidelines do not account for biological drivers of individual food preferences and consumption.

Dietary intake is significantly influenced by genetics, with the SNP heritability estimates for traits such as fruit, cheese, and alcohol intake around 10% [3]. Genome-wide association studies (GWAS) have identified hundreds of loci associated with eating behavior, from intake of specific foods to broader food groups, dietary patterns, and preferences [3,4].

Despite the growing body of evidence that the X chromosome harbors genetic variation associated with a wide range of complex traits [5–8], GWAS of diet have largely ignored the X chromosome. Historically, the X chromosome was omitted due to technical challenges in accurate genotyping and imputation introduced by haploid male genotypes and the pseudo-autosomal regions (PARs). However, now that robust X chromosome imputation methods have been available for over a decade [9] and have been implemented in large, commonly used data sets, including the UK Biobank (UKB), its omission persists largely due to unique analytical challenges arising from X chromosome inactivation (XCI) – the biological process that accounts for the difference in dosage of X-linked genes between diploid females and haploid males. XCI functions in females through the epigenetic silencing of one of the two copies of X in each cell early in fetal development. As cells proliferate, XCI in females creates a mosaic pattern of expression where, at each genomic position, one copy of X remains inactive, meaning cells express either the maternal or paternal allele depending on which X remains active. For many X-linked genes, the copy of X which is silenced is random and results in roughly equal expression of the two copies of X when cells are considered in aggregate across the body [10]. Under the assumption of random XCI, association analyses can assume that genotype dosage will be equivalent on-average between hemizygous males and homozygous females [11], and heterozygous females will have roughly equal levels of expression of the two copies of X. However, nearly one quarter of genes are believed to escape XCI to varying degrees, with additional variation between individuals and across tissue types [10].

Various models have been developed to statistically account for the complexities of XCI in X-chromosome-wide association tests (X-WAS). The most common model assumes random XCI in females [11] as described above, such that males are coded 0/2 and females 0/1/2. Association analysis can also be conducted in males and females separately, which avoids assumptions about dosage equivalence across sex, but substantially reduces statistical power, particularly in haploid males. Sex-combined models that include a SNP-sex interaction term better allow the genetic effect to vary by sex [12]. Additionally, Chen et al. 2021 propose an “X-factor” model for X chromosome association analysis, which includes both a SNP-sex interaction term and a heterozygosity term that captures non-additive effects in heterozygous females (termed dominance deviation). This statistically accounts for skewed XCI that could occur in heterozygous females.

Given the growing evidence that X chromosome variation contributes to complex traits, we hypothesize that the X chromosome harbors novel genetic variation associated with dietary intake. Here we apply a suite of X chromosome association models to maximize discovery of loci relevant to dietary intake in up to 445,773 individuals from multiple ancestries in the UKB.

## Results

### X-WAS models identify eighteen X chromosome loci associated with dietary intake

We performed X-WAS analysis on 46 dietary intake traits in up to 424,758 European (EUR) genetic ancestry individuals in the UKB using five distinct X chromosome models to account for XCI and maximize discovery (Table 1, Supplemental Tables S1-3). Traits included quantitative traits (e.g., fruit intake, poultry intake), binary traits (e.g., decaffeinated vs caffeinated coffee, reduced fat vs full fat milk), and principal component (PC) derived food groups (Supplemental Table S4). The five models included standard additive association tests in males-only, females-only, and in both sexes combined; an additive sex-combined model with a sex-interaction term; and the “X-factor” model, which included both sex-interaction and dominance deviation terms (see Methods). We also applied the standard additive model in five additional ancestry groups—African (AFR, N = 6709), Admixed American (AMR, N = 972), Central/South Asian (CSA, N = 8964), East Asian (EAS, N = 2766), and Middle Eastern (MID, N = 1604)—and performed cross-ancestry meta-analysis (up to N = 445773).

**Table 1.** Summary of X chromosome association models.

| MODEL | XCI ASSUMPTIONS | SOFTWARE | # SIGNIFICANT INDEPENDENT LOCI | STRENGTHS | LIMITATIONS |
| --- | --- | --- | --- | --- | --- |
| **Additive sex-combined** | Random | REGENIE | 12 | - Maintains full sex-combined sample size<br>- Compatible with common GWAS software<br>- High statistical power under random XCI | - Assumes male-female dosage equivalence<br>- Does not account for escape or skewed XCI<br>- Does not allow genetic effects to vary by sex |
| **Additive female-only** | Random or Escape | REGENIE | 2 | - Estimates female-specific genetic effects<br>- Avoids assumptions about male-female dosage equivalence | - Reduced sample size and limited power<br>- Unknown individual-level variability in XCI |
| **Additive male-only** | None | REGENIE | 5 | - Estimates male-specific genetic effect<br>- Avoids assumptions about male-female dosage equivalence and XCI | - Reduced sample size and limited power<br>- Fewer total alleles |
| **Sex-interaction** | Random or Escape | REGENIE | 10 | - Maintains full sex-combined sample size<br>- Allows genetic effect to vary by sex | - Unknown individual-level variability in XCI<br>- Does not account for skewed XCI |
| **X-factor** | Random, Escape, or Skewed | PLINK2 | 9 | - Maintains full sex-combined sample size<br>- Allows genetic effect to vary by sex<br>- Models heterozygosity effects in females<br>- Robust to underlying XCI status | - Limited implementation in mixed model software results in a reduction in sample size by removing related individuals<br>- Unknown individual-level variability in XCI |

Across all models, we identified 18 loci (Supplemental Figure S1A-R) associated with at least one of 20 dietary intake traits on the X chromosome (Table 2, Fig 1A), including 17 lead variants with no known associations in the GWAS Catalog [13]. Conditional analysis confirmed that two loci associated with adding salt to food 638KB apart (lead SNPs rs56157110 and rs6638366) are independent (rs6638366 *P* = 4.04 x 10^-11^ after conditioning on rs56157110, Supplemental Figure S2).

**Table 2.**
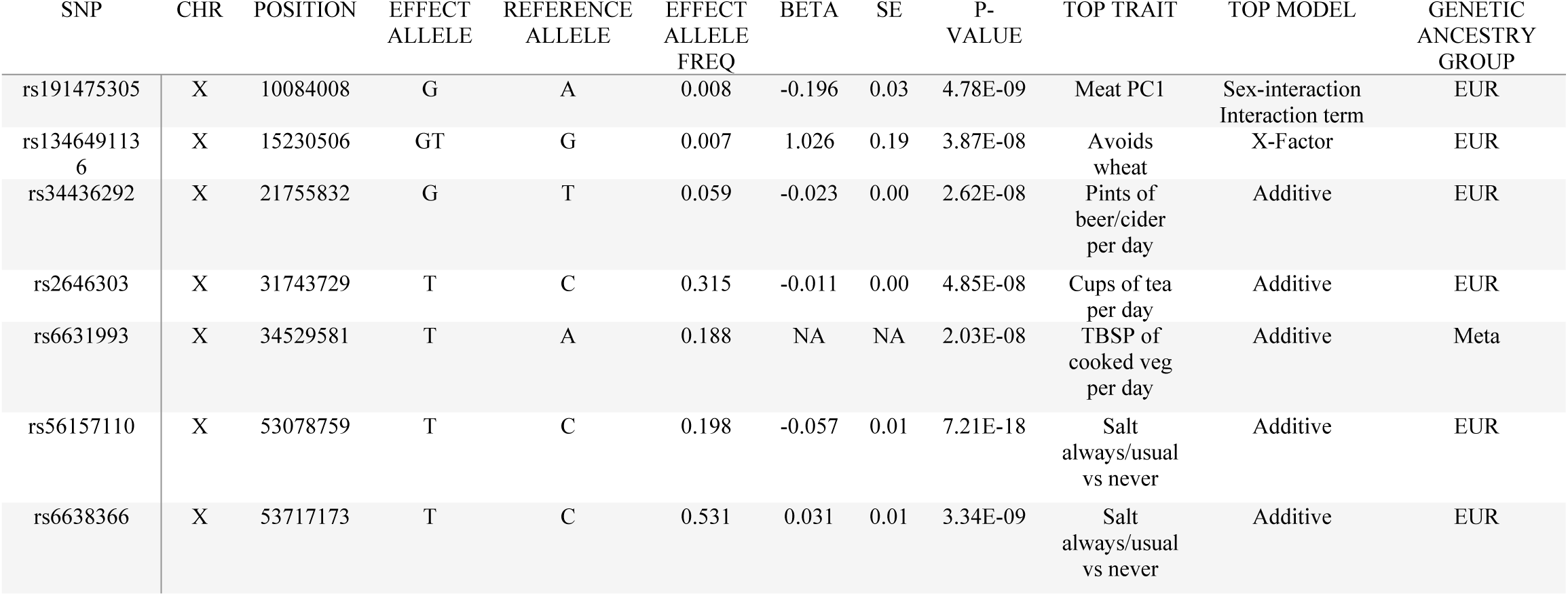
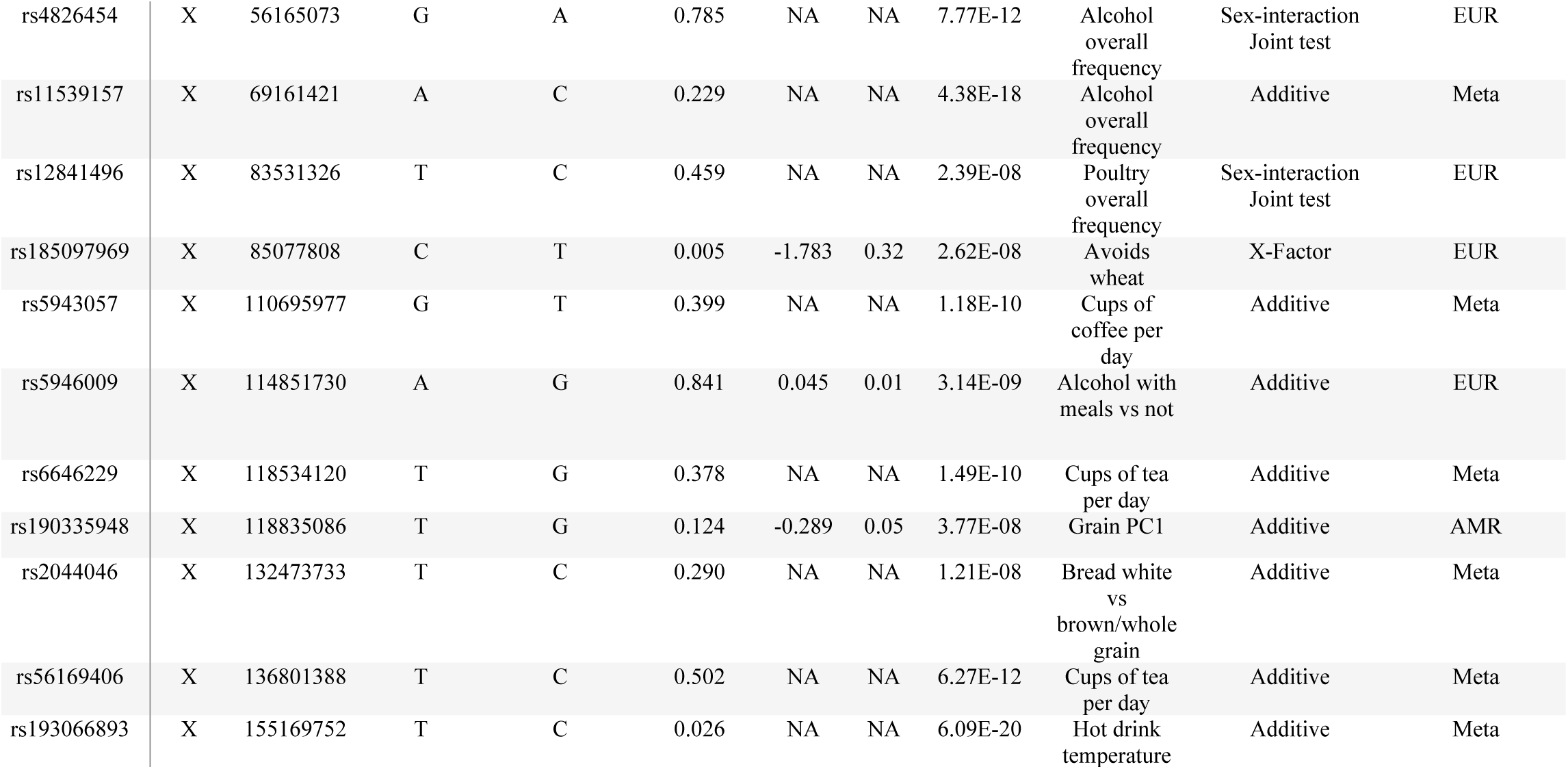
Eighteen loci significantly associated with dietary intake.

**Fig 1.**
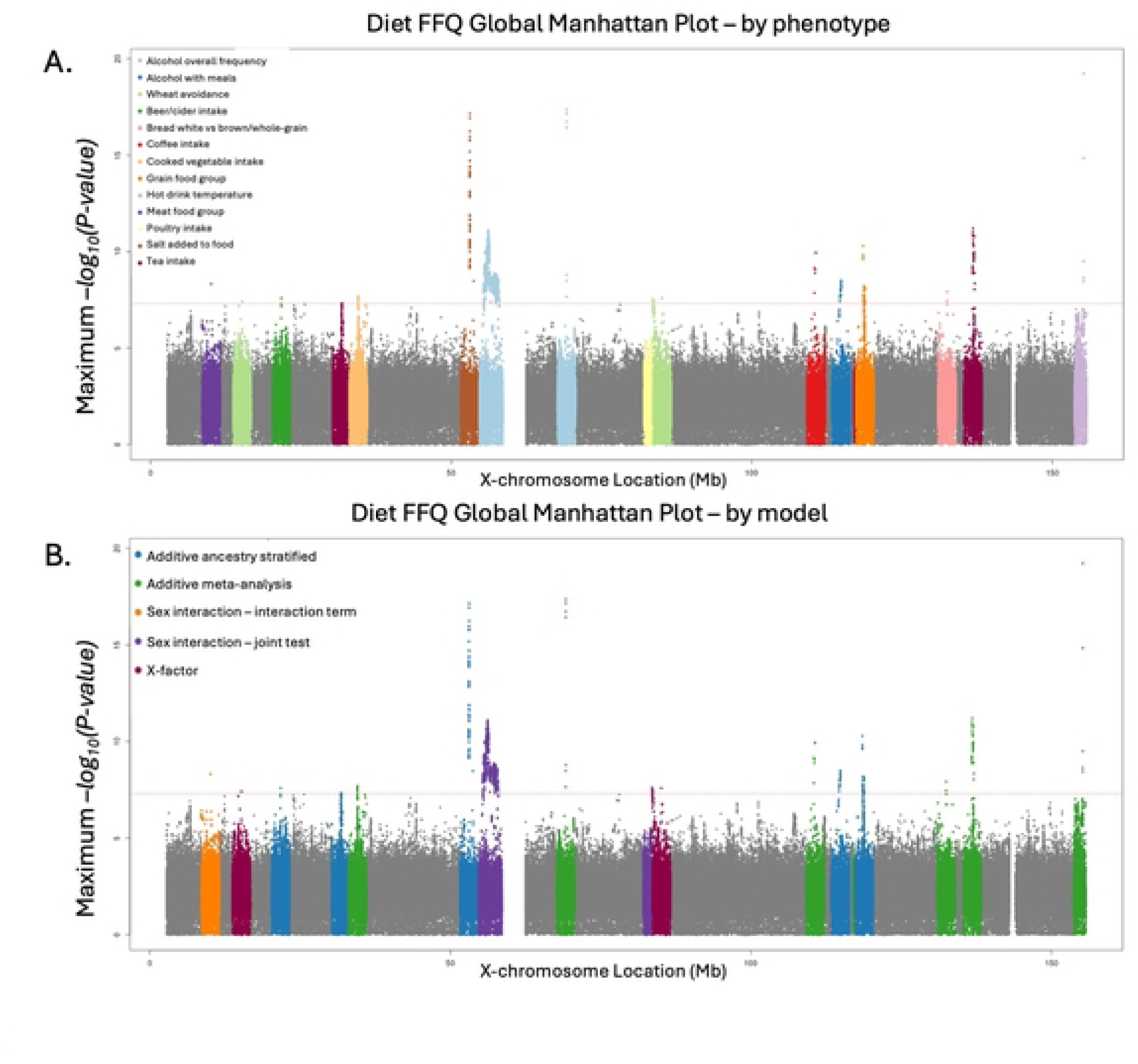
The most significant P-value for each SNP in an association test across all X-WAS models and multi-ancestry meta-analysis. Colors indicate A) the most significantly associated phenotype, or B) the model with the most significant association for any phenotype. Seven dietary intake phenotypes (cereal intake, bread intake, pork intake, processed meat intake, sugar avoidance, the drink food group PC, and the global dietary intake PC) had significantly associated variants but were not the most significantly associated phenotype for any of the genome-wide-significant loci. Similarly, significant variants were identified in the male-only and female-only association test models, but the sex-stratified models did not produce the most significant association for any genome-wide-significant loci.

Genome-wide significant associations were identified in all five tested models, however all male-only or female-only associations were more significant in the sex-combined additive model (Fig 1B, Supplemental Figure S3A-R). Five loci were more significant in the sex-interaction or X-factor model versus the standard sex-combined additive model, and three of these were only genome-wide significant in one of these more robust models, demonstrating their added value for X chromosome discovery. For seven loci, the most significant P-value was found in the cross-ancestry meta-analysis, suggesting that effects were broadly consistent across ancestry groups, and the increase in power from larger sample size outweighed any heterogeneity in effects. Of these, one locus, led by SNP rs6631993 associated with cooked vegetable intake (*P* = 2.03 x 10^-8^), was discovered only in the multi-ancestry meta-analysis. Five loci were most significant in the EUR-only additive sex-combined model, potentially suggesting some degree of heterogeneity in effects across ancestry groups and demonstrating our limited power to detect associations in groups with smaller sample sizes. One locus was specific to the AMR ancestry group: SNP rs190335948, ∼8KB downstream from the gene *ZCCHC12,* was associated with the grain food group PC (beta = -0.29, *P* = 3.77×10^-8^) and not associated in any other ancestry group (all *P* > 0.05).

We identified one significant sex interaction (Fig 2A). SNP rs191475305 is an intronic variant in the *WWC3* gene associated with the meat food group PC (beta = -0.20, p = 4.78×10^-9^) with opposite effects on intake in females versus males (Fig 2B). It was therefore not significant in the sex-combined additive model without the sex-interaction term (Figure 2C) and was nominally significant in females-only (*P* = 3.56 x 10^-7^) and males-only (*P* = 0.0057). *WWC3* has not previously been linked to eating behavior, but does contain regions of sex-differential methylation patterns associated with tobacco smoking behavior [14]. Another locus led by intergenic SNP rs12841496, 19KB downstream of *POU3F4*, was significantly associated with poultry intake in the additive sex-combined model (beta = 0.0098, *P* = 4.67×10^-8^), but the association was more significant when the model included a sex interaction term (*P* = 2.39×10^-8^), suggesting that although the interaction itself did not reach genome-wide significance, accounting for it reduced heterogeneity.

**Fig 2.**
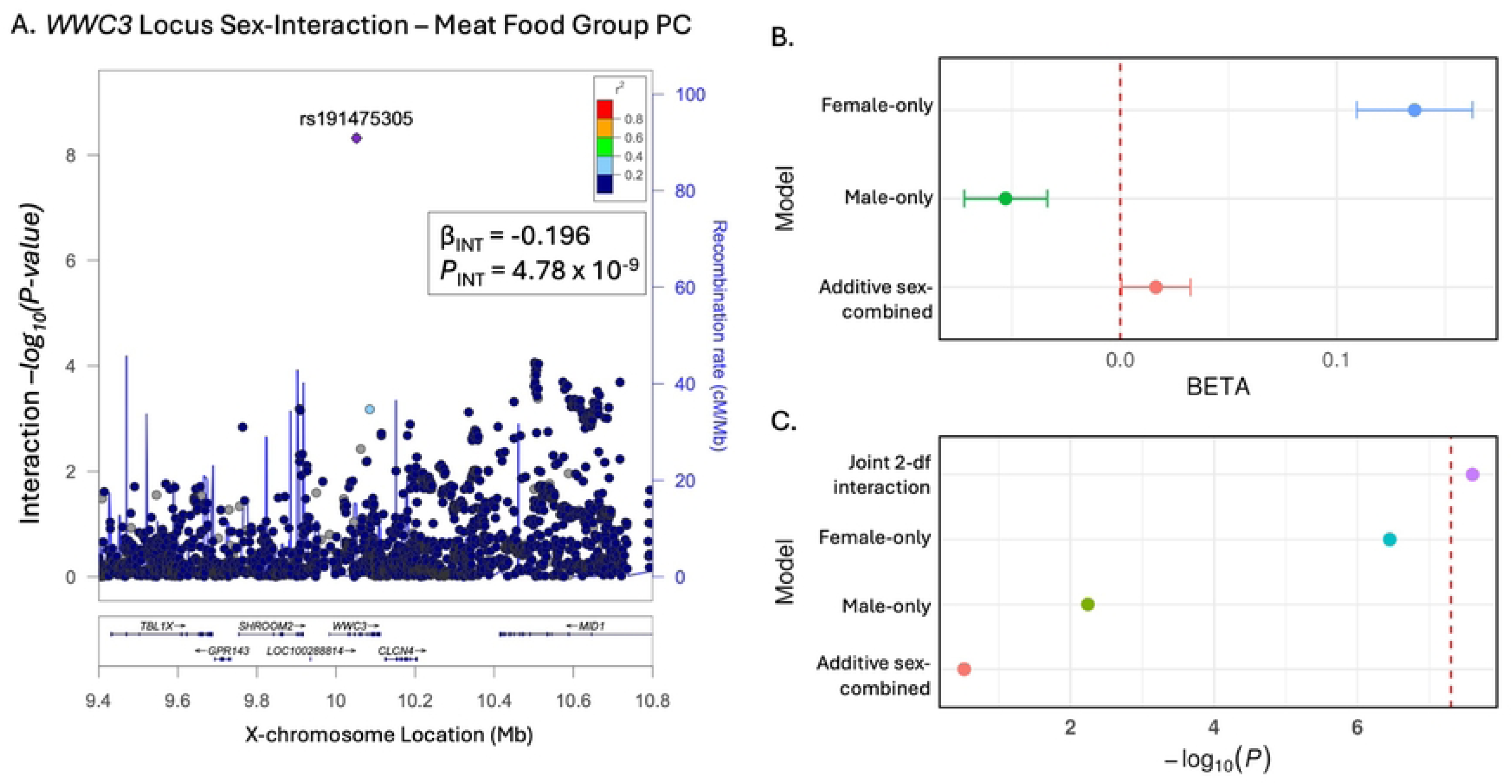
We identified one significant sex interaction: rs191475305 associated with the meat food group PC. A) Interaction term p-values for the locus around *WWC3* with a significant sex-interaction associated with the meat food group PC (beta = -0.196, *P* = 4.78 x 10^-9^); B) P-values of the top SNP (rs191475305) for the meat food group PC in different X-chromosome models (Joint 2-df interaction refers to the joint 2-degree-of-freedom interaction test in REGENIE); C) effect sizes of the top SNP (rs191475305) for the meat food group PC in different X-chromosome models (the joint interaction test does not have an effect size, only a p-value, so it is not included in panel C).

Furthermore, SNP rs56157110, located in the 3’ UTR of *GPR173*, was associated with adding salt to food. The strongest association was observed in the additive sex-combined model (beta = -0.057, *P* = 7.21×10^-18^) and remained genome-wide significant in the male-only model (beta = -0.060, *P* = 1.46×10^-13^), but not the female-only model (beta = -0.049, *P* = 1.32×10^-5^). Although the effects were similar between males and females, the standard error was higher in females (SE_females_ = 0.011 vs SE_males_ = 0.008), consistent with differences in effective allele dosage or additional heterogeneity in females due to XCI (Supplemental Figure S2F). Additionally, rs56157110 is an eQTL for *TSPYL2* in coronary artery tissue which is associated with expression in males (*P* = 0.002), but not in females [15].

Several additional loci showed a similar pattern between males and females. SNPs rs5946009, an intronic variant within *HTR2C*, associated with alcohol consumption with meals; rs2044046, intronic variant within *MBNL3*, associated with white vs brown or whole grain bread; rs56169406, an intergenic variant near *ARHGEF*, associated with tea intake; and rs11539157, a missense variant in *PJA1*, associated with alcohol intake were all most significant in the additive sex-combined model, but remained genome-wide significant in the male-only model. Effect estimates were similar but attenuated in the female-only analysis accompanied by larger standard errors (Supplemental Figures S2M, S2P, S2Q, S2I).

We identified two independent loci 13MB apart, both associated with overall alcohol intake in the sex-combined additive model (Fig 3A). In addition to the locus led by rs11539157, a missense variant in *PJA1* (Fig 3B, Supplemental Figure 3I), we identified a locus led by rs4826454 that overlaps multiple genes (Fig 3C, Supplemental Figure 3H) including *FOXR2*. Previous research has shown that *FOXR2* and *PJA1* interact, with *PJA1* regulating apoptosis through degradation of the fork-head box R2 (FOXR2) protein [16]. We find that they each have sex-specific effects on alcohol intake (Figure 3D), with rs4826454 being most significant in the sex-interaction joint test (*P* = 7.77 x 10^-12^) but remaining significant in the female-only model (beta = 0.024, *P* = 2.41 x 10^-11^). The SNP-sex interaction term was marginally significant for rs4826454 (*P* = 5.05 x 10^-5^).

**Fig 3.**
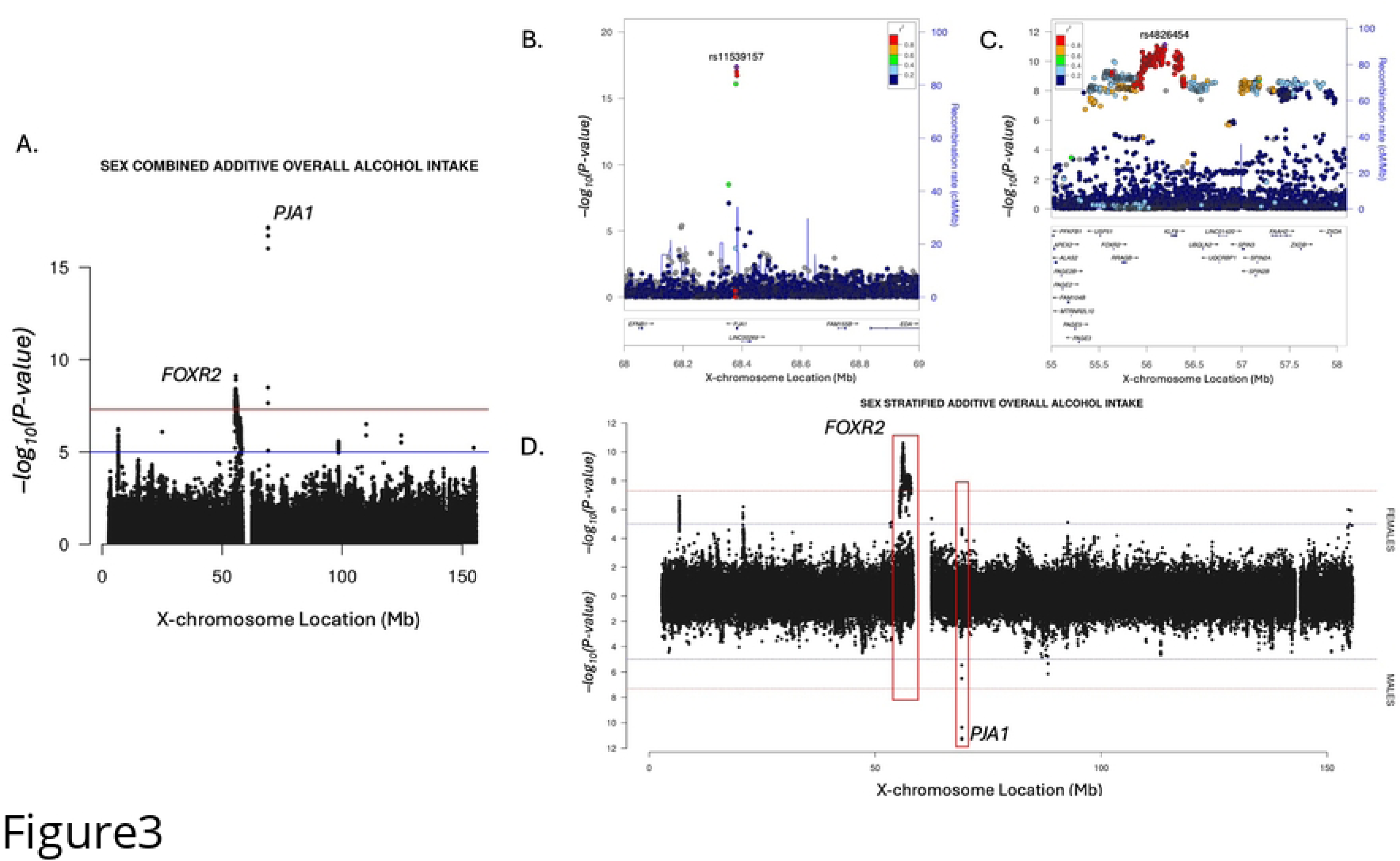
A) Two loci significantly associated with overall alcohol intake around *FOXR2* and *PJA1* genes in the sex-combined additive model. B) Locus led by rs11539157 was most significant in the sex-combined meta-analysis. C) Locus led by rs4826454 was most significant in the sex-interaction model. D) Each contains sex-specific signals related to alcohol intake.

Two loci, led by rs1346491136 (intergenic, 13.5KB downstream of *ASB9*) and rs185097969 (intronic within *APOOL)*, were significantly associated with wheat avoidance only in the X-factor model. This suggests genetic effects that deviate from XCI assumptions in the standard additive model, potentially due to non-additive effects in heterozygous females. SNP rs1346491136 was also nominally significant in the female-only model (*P* = 0.03), providing some evidence for an association in females, although modest.

### Many loci may be associated with dietary intake through more complex pathways

Given dietary intake represents a series of complex behavioral traits correlated with many diseases and markers of health, social determinants of health, and other lifestyle factors, we sought to better characterize the complete phenotypic spectrum associated with each of our top loci through phenome-wide associations studies (PheWAS) using X-WAS data from 924 traits in Pan-UKB (up to N = 420,531). Nine out of 17 tested loci had genome-wide significant associations with non-diet traits (Table 3), primarily biomarkers and anthropometric measurements (Supplemental Table S5). Colocalization analyses between association signals for diet and top non-diet PheWAS traits identified five loci (Supplemental Figure S4A-E) with strong evidence of colocalization (PP4 > 0.5), indicating that the diet and non-diet associations at these loci are likely driven by the same underlying causal variant.

**Table 3.** PheWAS Summary.

| SNP | TOP DIETARY INTAKE TRAIT | TOP PHEWAS TRAIT | TOP PHEWAS P-VALUE | COLOC PP H4 | # GWS PHEWAS TRAITS |
| --- | --- | --- | --- | --- | --- |
| rs56157110 | Salt always/usual vs never | Insulin-like growth factor 1 | 3.98E-49 | 0.992 | 10 |
| rs6638366 | Salt always/usual vs never | Body mass index | 3.39E-26 | 2.41x10<sup>-9</sup> | 39 |
| rs4826454 | Alcohol overall frequency | Mean platelet volume | 1.26E-218 | 0.0996 | 44 |
| rs11539157 | Alcohol overall frequency | Impedance of whole body | 5.75E-64 | 0.998 | 46 |
| rs5943057 | Cups of coffee per day | Sex hormone-binding globulin | 3.98E-430 | 0.993 | 75 |
| rs6646229 | Cups of tea per day | Gamma glutamyl transferase | 6.92E-22 | 0.481 | 10 |
| rs2044046 | Bread white vs brown/wholegrain | Impedance of leg (right) | 5.89E-13 | 0.02 | 12 |
| rs56169406 | Cups of tea day | Body mass index | 3.47E-14 | 0.944 | 22 |
| rs193066893 | Hot drink temperature | Leg predicted mass (right) | 2.61E-10 | 0.998 | 9 |

*GWS = genome-wide significant*

In contrast, four dietary intake loci (Supplemental Figure S4F-I) did not colocalize with their top non-diet PheWAS associated traits, providing evidence that these signals are driven by distinct causal variants. This includes rs6638366 for adding salt to food and BMI (probability for distinct causal variants [PP3] = 0.99), rs4826454 for overall alcohol intake and mean platelet volume (PP3 = 0.90), and rs2044046 for white vs brown or whole grain bread intake and impedance of the leg (PP3 = 0.96).

Finally, we conducted sensitivity analysis at each lead dietary association, adjusting for several covariates suspected as confounders (Supplemental Table S6). After including 19 additional covariates, including derived socioeconomic status, BMI, total energy intake, and disease diagnoses, we found two SNPs with greater than 10% attenuation in both effect size (Supplemental Figure S5) and significance (Supplemental Figure S6). SNP rs5943057 associated with coffee intake was greatly attenuated after adjustment for additional covariates (beta_baseline_ = 0.01, *P_baseline_* = 0.0011, beta_sensitivity_ = 0.0088, *P_sensitivity_* = 0.0059), and rs34436292 associated with beer/cider intake was largely affected by adjustment for additional covariates (beta_baseline_ = - 0.012, *P_baseline_*= 0.11, beta_sensitivity_ = -0.0097, *P_sensitivity_* = 0.17). Although the SNP effects were no longer GWS with the reduced sample size, the attenuation of effects when accounting for potential confounders suggests that these two observed SNP-diet associations may be largely attributable to relationships with non-diet traits, rather than being true diet-specific genetic effects.

## Discussion

We identified 18 loci on the X chromosome associated with 20 dietary intake traits in the UKB using multiple association models that account for different XCI assumptions. These findings add to the growing evidence that the X chromosome harbors genetic variation with important effects on a wide range of phenotypes, including complex behavioral traits like dietary intake.

Most loci reached genome-wide significance in the standard sex-combined additive association model, which demonstrates that although XCI-robust models can improve signal and identify novel loci, the greatest benefit may come from simply including the X chromosome in GWAS analyses. The ability of commonly used GWAS software such as PLINK [17], REGENIE [18], and SAIGE [19] to appropriately model male hemizygosity and female diploidy under the standard additive framework makes routine inclusion of the X chromosome in GWAS analyses straightforward and should improve the landscape of X chromosome associations moving forward.

Nevertheless, more complex X chromosome models allowed us to test associations with dietary intake when the assumption of additivity does not hold at the population level and to investigate SNP-sex relationships affecting dietary intake. Three associations were identified only with these models, including one significant SNP-sex interaction associated with meat intake and two loci associated with wheat avoidance detected using the “X-factor” model. Of note, two loci that were GWS only in the X-factor model were identified in an unrelated subset of individuals in PLINK, were not significant in the REGENIE sex-combined additive model or the sex interaction models (*P* > 0.05), and are relatively rare (MAF ∼ 0.005), suggesting these signals should be interpreted cautiously and require replication. This highlights the potential value of implementing the X-factor model’s dominance deviation term within the current gold-standard GWAS software that better account for relatedness and population structure.

Several of our dietary intake associations are supported by functional evidence in the literature. SNP rs5946009, an intronic variant within *HTR2C*, was associated with whether alcohol was typically consumed with meals. *HTR2C* encodes a serotonin receptor protein which is known to regulate appetite and eating behavior [20,21], as well as response to stress [22]. SNP rs5946009 was not genome-wide significant for other phenotypes in the PheWAS, and in our covariate sensitivity analyses, the effect was only modestly attenuated (<10%), with socioeconomic status (SES) variables being the strongest predictors. SNP rs56157110, located within the 3’ UTR of *GPR173* and acting as an eQTL for *TSPYL2* in multiple tissues is associated with adding salt to food. A recent study by Haddock et al. identified an interaction between *GPR173* and *PNX* regulating thirst behavior and hypothesized that these genes could be important to maintaining fluid and electrolyte balance [23].

Interpretating genetic effects on complex, behavioral phenotypes like dietary intake and distinguishing true signal from confounding effects is challenging because many diet-associated variants also show associations with non-diet traits. Several of our loci had extremely significant associations with non-diet traits, including anthropometric traits and biomarkers. Colocalization analysis between these signals and our dietary intake signals support common underlying causal variants at some loci. This pattern reflects the complex relationships between dietary intake and metabolic health. It is difficult to statistically determine the true causal mechanism underlying associations, particularly whether a variant has pleiotropic effects, whether genetic influences on health alter eating behavior or vice versa, or whether other environmental or social factors are confounding observed relationships.

There are some important limitations to this study. Although the use of multiple X chromosome association models allowed us to explore different statistical assumptions about XCI status at the population level, patterns of expression vary between individuals and between tissues within an individual. True XCI status can only be determined through integration of individual-level expression data. Additionally, dietary intake phenotypes were derived from brief food frequency questionnaires in the UKB, which are subject to self-reporting biases and capture only a limited number of food groups, rather than specific food items, which may have unique genetic determinants.

In summary, our X-WAS identified 18 loci associated with a range of dietary intake phenotypes, including both individual food items and broader food groups, using multiple association tests that account for XCI and SNP-sex interactions. Together these results demonstrate that the X chromosome harbors genetic variation influencing even complex, behavioral traits like dietary intake. The ability to model random XCI in widely used GWAS software should facilitate broader inclusion of the X chromosome in future genomic association studies.

## Methods

### Dietary Intake Phenotypes in the UKB

We curated 46 dietary intake traits from the food frequency questionnaire (FFQ) in the UKB (Supplemental Table S4), within six different ancestry groups: up to 424,758 EUR; 6,709 AFR; 972 AMR; 8,964 CSA; 2,766 EAS; and 1,604 MID. We defined 22 quantitative and 17 binary traits following the methods for the autosomal GWAS of dietary intake by Cole et al. [3] and seven food group PCs following Westerman et al. [24].

Quantitative traits were all converted to intake per day and averaged across all available FFQs for each individual. Specific alcohol phenotypes (e.g., red wine, beer/cider, etc.) were derived from weekly or monthly alcohol intake fields, depending on their response to overall alcohol intake frequency (field 1558). Over 90% of responses to intake of fortified wine and “other alcoholic drinks” were zero and were therefore dropped from analysis. Spirit intake was converted to a binary consumption variable because over 60% of responses were zero. Three ordinal phenotypes (adding salt to food, preferred hot drink temperature, and fat content of milk consumed) were converted to a continuous scale. All quantitative traits were adjusted for covariates and inverse rank normal transformed (IRNT) within genetic ancestry groups as defined by the Pan-UKB [25]. Covariates included: sex (field 31), average age across FFQs, age^2^, age*sex, age^2^*sex, baseline assessment centre (field 54), birthplace (field 1647), genotyping array (field 22000), and number of FFQs taken. The first 20 genetic PCs were included as covariates directly in the GWAS. Binary traits were derived from baseline FFQ responses only, with all covariates adjusted for directly in the GWAS.

Food group PCs were the first PC from a principal component analysis (PCA) on groups of related foods using pre-adjusted, IRNT quantitative traits in unrelated individuals, as follows: 1) fruits and vegetables, 2) non-alcoholic drinks, 3) fish, 4) meat, and 5) grains. Missing values were median imputed prior to PCA. For the fruits and vegetables, fish, and meat food groups, the first PC represented more vs less intake of items in that food group (Supplemental Figure S7A-C). The first PC of the “drinks” food group represented more tea intake vs more coffee intake (Supplemental Figure S7D), and the first “grain” PC represented more cereal intake vs more bread intake (Supplemental Figure S7E). We also included the second PC from the fruits and vegetables PCA, as it more closely represents more fruit intake vs more vegetable intake. We created a global dietary pattern phenotype by including all the individual foods included in any food group together in PCA, plus additional overall alcohol intake and cheese intake. The first global PC differentiates more fish, vegetable, and fruit intake vs more meat, grain, and cheese intake, representing a general healthful dietary pattern, as has been observed previously in these types of analyses on FFQ data [3,26] (Supplemental Figure S7F).

### UKB Genetic Data and X Chromosome-Specific Quality Control

Our analyses utilized UKB genetic data imputed to the TOPMed reference panel [27] and accessed through the UKB Research Analysis Platform under UKB application 100578. Methods for genotyping, imputation, and initial quality control (QC) procedures have been described previously [28,29].

We applied a series of QC steps to the X chromosome adapted from existing recommendations [30–32], applied within Pan-UKB genetic ancestry groups using PLINK v2.00a3.1. The PARs were removed, since they are small, contain few genes, and can be analyzed without the need for additional X-specific considerations. We also removed samples with sex mismatches (using PLINKv1.90b6.26 --check-sex function) and high missingness across genotypes (>5%). We applied variant filters within females, within males, and then all together to remove variants that: 1) were out of Hardy-Weinburg Equilibrium in females (p < 1×10^-6^); 2) had high missingness across individuals (>2%) in males, females, or all together; 3) had fewer than 25 counts of the minor allele within males or within females. This resulted in a subset of 425,081 samples (195,327 males and 229,754 females) and 1,332,895 variants for use in association testing in the largest ancestry group (EUR).

### X-wide Association Models

We applied five different X chromosome association models (Table 1) in the largest ancestry group in the UKB (EUR, N up to 424,758) to maximize discovery and investigate whether significant associations were robust to differing assumptions about XCI.

First, we fit a standard additive association model on all individuals (equation 1), which assumes each additional copy of the effect allele contributes to an equal change in the phenotype. This model assumes random XCI by coding female genotypes as 0/1/2 and male genotypes as 0/2. We also fit the standard additive association model in females-only (assuming random or escape XCI) and males-only (where XCI is irrelevant); although these models remove any noise due to heterogeneous effects across the sexes, enabling the discovery of sex-specific effects, they also dramatically reduce sample size.

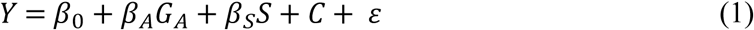

*Y* is the phenotype, *β_A_* is the additive genetic effect, *β_S_* is the effect of sex, and *C* represents additional covariates.

To consider sex-specific differences in effects while retaining statistical power, we fit a model with a SNP-sex interaction term (G*S, equation 2) and evaluated the significance of both the interaction term itself and the two degrees-of-freedom joint test accounting for the SNP-sex interaction. All the above models were tested in REGENIE (v3.1.1).

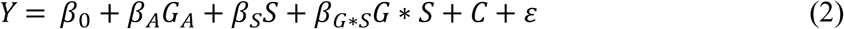

Additionally, we tested the “X-factor” model (equation 3) proposed by Chen et al. [12], which includes a SNP-sex interaction term and a dominance deviation term (*G_D_*), which is coded 0/1 for heterozygosity at each variant (one for heterozygous females and zero for homozygous females and hemizygous males). This term allows for deviation from additivity, which might arise under non-random patterns of XCI, particularly in the case of skewed XCI where heterozygous females might have unequal expression of the two copies of X. REGENIE does not support this model; therefore, we applied this model in 361,009 unrelated EUR individuals in PLINK2.

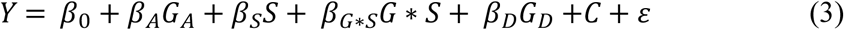

We repeated the additive sex-combined association model on five additional ancestry groups as defined by Pan-UKB [25]. X-specific QC filters and phenotype derivation was conducted within each ancestry group. Multi-ancestry meta-analysis was performed to maximize power, using the sample-size weighted method in METAL [33] and a total of 445,773 participants.

### Classifying independent signals

GWAS were filtered to remove rare variants (MAF < 0.005) and variants with low imputation quality (INFO < 0.3). Independent loci were defined based on distance using a large 3MB window around the lead SNP surpassing genome-wide significance (*P* ≤ 5.0 x 10^-8^). For loci that appeared on the regional Manhattan plot (LocusZoom) to have a secondary signal, we repeated the association test conditioning on the lead variant, iterating this process by adding the secondary signal’s lead variant to the model until no variants remained genome-wide significant. Conditional analysis also confirmed that two nearby loci separated by more than 3MB were not independent and therefore were collapsed into one (Supplemental Figure 1H).

### Characterizing associations with other traits at each top locus

To gain a clearer understanding of the complete spectrum of phenotypic associations for 17 of our top loci (rs1346491136 was not in Pan-UKB), we conducted PheWAS and colocalization analysis. After performing LiftOver [34] of our top variants to GRCh37, we performed a PheWAS at each lead variant using public GWAS from Pan-UKB project, covering 924 traits in the UKB that passed QC requirements in the EUR group (or AMR group for rs190335948), as described by Pan-UKB.

For each locus with significant (*P* < 5.0 x 10^-8^) associations with one or more non-diet-related traits in Pan-UKB, we performed colocalization analysis to estimate the posterior probability that the two signals shared an underlying causal variant using the coloc (v5.1.0.1) R package [35]. We used the EUR Pan-UKB results for all variants except rs190335948, which was AMR specific. We considered posterior probabilities for a shared causal variant greater than 0.5 to be colocalized signals.

To evaluate the influence of confounding factors on the genetic association with dietary intake, we performed additional covariate sensitivity analyses for our top SNPs. For each lead SNP, we compared SNP effect and significance between a baseline model that matched our original GWAS test and an adjusted model that included 19 additional covariates in 122,943 unrelated EUR individuals with complete covariate information in R (v4.4.0) [36]. Covariates included: 1) self-reported major dietary changes (field 1538); 2) self-reported diagnosis of diabetes (field 2443), removing those with only gestational diabetes (field 4041); 3) self-reported diagnosis of heart attack (field 6150); 4) average BMI across visits, inverse-normal transformed (field 21001); 5) binary obesity based on BMI >30; 6) self-reported blood pressure medication use (fields 6177 and 6153); 7) cholesterol medication use (fields 6177 and 6153); 8) continuous triglyceride levels (field 30870); 9) binary very high triglycerides defined as >5.65mmol/L; 10) self-reported non-white ethnicity (field 21000); 11) self-reported pregnancy at baseline (field 3140); 12) self-reported depression at baseline (field 20126); 13) cancer diagnosis within 1yr of taking an FFQ; 14) physical activity (field 22039) at baseline; 15) derived total energy intake averaged across instances (from field 26002); and 16-19) derived SES, including household income (from field 738), educational attainment (from field 6138), employment status (from field 6142), and a composite SES variable. For the composite SES variable, individuals with missing data for two or more of these SES variables (income, education, employment) or with “Prefer not to answer” responses for all three variables were excluded. For the remaining participants, the mice (v 3.17.0) R package [37] was used to impute values for any non-responses (NA; “Prefer not to answer”; “Do not know”) across the three SES variables using age, sex, assessment center, and Townsend deprivation index as additional predictors. Latent class analysis (m=5, maxit=10) was performed with the poLCA (v 1.0.6.1) R package [38] to identify latent individual-level SES groups for all five imputed datasets. The modal class assignment was used for the final composite SES categorical variable. Likewise, the modal imputed value was assigned for any non-responses for the final individual SES variables.

## Acknowledgements

We are deeply indebted to the participants in the UK Biobank, without whom this work would not be possible. This research has been conducted using the UK Biobank Resource under application number 100578. This work was supported by the Alpine HPC system, which is jointly funded by the University of Colorado Boulder, the University of Colorado Anschutz, Colorado State University, and the National Science Foundation (award 2201538).

## Data Availability Statement

X-WAS summary statistics will be made available through the GWAS catalog (accession number to-be-provided).

## Author Contributions Statement

JBC and MSB designed the study. Analyses were performed by MSB. KJS, JBC, and MSB curated dietary phenotypes. WBP curated SES variables. The primary draft was written by MSB and JBC, with input and approval from KJS and WBP.

## Supporting Information Captions

Table S1. Genome-wide significant associations from all models using REGENIE

Table S2. Genome-wide significant associations from X-factor model (PLINK2)

Table S3. Genome-wide significant associations from multi-ancestry meta-analysis (METAL)

Table S4. Dietary intake phenotypes evaluated in the UK Biobank

Table S5. Top SNPs associations with non-diet traits in PheWAS

Table S6. Effects of potentially confounding covariates

Figure S1A-R. Regional Manhattan (LocusZoom) plots for diet loci

Figure S2. LocusZoom plot of two independent signals associated with salt intake within 3MB

Figure S3A-R. Forest plots of SNP effects and significance across X-WAS models

Figure S4A-I. Colocalization of diet-identified signals with associated non-diet traits

Figure S5. Covariate sensitivity changes in SNP effects

Figure S6. Covariate sensitivity changes in p-values

Figure S7A-F. Food group principal components

